# Abdominal-B regulates the male seminal fluid transferome and new female fecundity factors required for sperm sex peptide binding

**DOI:** 10.64898/2026.01.10.698609

**Authors:** Kathrin Steck, Lina Verbakel, Peter Kerwin, Vladimir Trajanovikj, Katarzyna Kjøge, Jan J. Enghild, Anne C. von Philipsborn

## Abstract

Seminal fluid proteins determine reproductive success in a wide range of animals. In the Drosophila male accessory gland, seminal fluid is mainly produced by two cell types, a vast majority of main cells and a small number of secondary cells that possess a specialized secretory apparatus with unusually enlarged dense core granule vesicles. Loss of Abdominal-B expression from secondary cells in the enhancer mutant *iab-6^cocu^* disrupts their transcriptional and secretory identity. Consequently, mutant males fail to induce the long-term post-mating response in females, which is characterized by a loss of receptivity and sustained egg laying. Here, we determine how secondary cells shape the seminal transferome and the female response by assessing *iab-6^cocu^* male accessory gland and female mate reproductive tract proteomes. We find downregulation of seminal fluid proteins that constitute a signaling network that enables sperm binding and the sustained action of the key regulator Sex peptide and identify two new Sex peptide network proteins crucial for female fecundity, Cornutus (CG1701) and Hanrej (CG42564). Cornutus is required for mating dependent dense core granule vesicle release, providing a link between the products of these compartments and the female long-term post-mating response. Our data highlights the importance of secondary cell signaling and secretion for overall seminal fluid composition and Sex peptide network function as well as the interdependence of main and secondary cells and their secretory products, advancing the general understanding of how seminal fluid signaling pathways modulate female physiology, sperm use and offspring production.

## Introduction

Seminal fluid is a key determinant for fertility and fecundity in internally fertilizing animals. It is not only a vehicle for gamete transport but also a “socially transferred material” (Hakala et al. 2023) fullfilling important signaling functions that affect both sperm and the female body. Seminal fluid is produced by secretory tissues of the male reproductive tract. It contains a multitude of proteins and peptides as well as other components, including nutrients, hormones, antimicrobials, pheromones, and lipid vesicles such as exosomes. In many animals, seminal fluid proteins (SFPs) mediate sperm survival and storage, influence ovulation and fertilization, and modulate female behavior and offspring health (Perry et al. 2013, Schjenken and Robertson 2020, Vallet-Buisan et al. 2023). In most species, the mechanisms by which SFPs exert their multifaceted effects are poorly understood and not readily accessible experimentally. *Drosophila melanogaster* has arguably the best studied seminal fluid proteome among genetic model organisms (Wigby et al. 2020). It has been instrumental for studying the effects of SFPs on females (Chapman et al. 1995, Kubli 2003, White and Wolfner 2022), postcopulatory sexual selection (Sirot and Wolfner 2015), the evolution of SFPs (McGeary and Findlay 2019, Sirot 2019, McQuarrie et al. 2025, Ranz et al. 2025), and male allocation (Wigby et al. 2009, Bretman et al. 2011, Hopkins et al. 2019a, Kerwin and von Philipsborn 2020). Currently, around 300 proteins are considered high confidence SFPs that are transferred from male to female. They fall into similar functional classes as the larger human seminal plasma proteome, such as proteases, protease inhibitors, redox-related proteins, proteins with immune function and proteins involved in lipid metabolism (Wigby et al. 2020, Hurtado et al. 2022).

Despite the complex and critical roles of seminal fluid in reproductive success, only a small fraction of individual Drosophila SFPs have been functionally characterized. We lack a comprehensive understanding of the signaling pathways involved in SFP production and allocation, as well as SFP interactions with the female reproductive tract. Most research on how SFPs influence female physiology and behavior has focused on Sex peptide (SP), which binds a neuronally expressed receptor in females and orchestrates post-mating changes in female receptivity, oogenesis and egg laying, feeding preferences and gut homeostasis, aggression and sleep (reviewed in Ellendersen and von Philipsborn 2017, Hoshino and Niwa 2021, Hopkins and Perry 2022). SP can only exert its sustained and wide-ranging effect when bound and gradually released from sperm in the female sperm storage organs (Peng et al. 2005). A conserved cofactor network of other SFPs is required for stable SP sperm binding and the long-term female post-mating response. The dynamics, regulation and extent of this so-called SP network are not yet fully understood (Singh et al., 2018, Carlisle et al. 2025).

Drosophila seminal fluid is mainly produced by the paired accessory glands (AGs). The Drosophila AG is functionally analogous to the mammalian prostate (Wilson et al. 2017), with which it shares developmental programs (Kumari and Sinha 2021), transcriptional networks (Church et al. 2025) and growth and secretory pathways (Corrigan et al. 2014, Leiblich et al. 2019, Sekhar et al. 2023). It consists of two secretory cell types, with around 1000 hexagonal shaped main cells and around 45 large, rounded secondary cells at the distal tip of the gland. Main and secondary cells have different transcriptional profiles. While SP is predominantly produced by main cells, expression of some SP network members (*lectin46-Ca*, *lectin46-Cb* and *CG17575)* is highly enriched in secondary cells. Secondary cells are characterized by a specialized secretory apparatus with unusually large vesicles containing exosomes and dense core granules (DCGs) (Bairati 1968, Avila et al. 2016, Prince et al. 2019, Immarigeon et al. 2021, Majane et al. 2022, Wells et al. 2023). For fully functional seminal fluid, interaction between both cell types and their products is required (Hopkins et al. 2019b). This is illustrated by the drastic effect of transcriptional dysregulation of secondary cells in a *Hox* gene enhancer mutant. Specific loss of Abdominal-B (Abd-B) expression in secondary cells of *iab-6^cocu^* mutant males transforms their typical morphology and abolishes the large DCG vesicles. Females mated to *iab-6^cocu^*mutant males fail to maintain the long-term post-mating response, show a high rate of remating and low fecundity (Iampietro et al. 2010, Gligorov et al. 2013) and do not respond with normal acoustic signals to seminal fluid in copula (Kerwin et al. 2020). SP and the main known SP network proteins were still found to be present in *iab-6^cocu^* mutant males (Gligorov et al. 2013). Transcriptome analysis of *iab-6^cocu^* mutant AGs, with functional analysis of selected candidates, indicated that most of the downregulated genes influencing the female long-term post-mating response were not SFPs but acted indirectly on seminal fluid composition by altering secondary cell physiology and secretory activity (Sitnik et al. 2016). The complex phenotype of *iab-6^cocu^*mutants, the contribution of secondary cell specific secretions to seminal fluid composition and a functional SP network are not yet fully understood.

To gain better insight into how the seminal fluid proteome and transferome are changed upon loss of Abd-B expression in secondary cells, we assessed protein abundances by mass spectrometry of male and mated female tissues. Among SFPs downregulated in the *iab-6^cocu^* mutant transferome, we found four previously characterized SP network proteins and discovered two additional SFPs that are indispensable for the long-term post-mating response. The two new fecundity and receptivity factors, *cornutus* (CG1701) and *hanrej* (CG42564), are required in secondary and main cells respectively, to suppress female remating and sustain ovulation by mediating SP sperm binding. Cornutus is required for mating induced DCG release and affects aspects of DCG maturation. Our findings highlight how specialized secretion pathways maintained by Abd-B expression in secondary cells shape the overall transferome from the AG by broadly regulating abundances of SP network proteins.

## Results

### Seminal fluid proteome and transferome changes in *iab-6^cocu^* mutant males

To understand how the seminal fluid proteome in the male accessory gland (AG) as well as the transferome reaching the female reproductive tract is changed in *iab-6^cocu^* mutants, we performed liquid chromatography tandem mass spectrometry of three tissue types: virgin male accessory glands (AG proteome non mated males), accessory glands of males collected immediately after the end of copulation (AG proteome mated males) and the female reproductive tract (RT) including uterus and sperm storage organs collected immediately after copulation (RT proteome mated female). For unmated and mated AG proteome assessment, we compared *iab-6^cocu^* mutants to wild type controls. For mated female RT proteome assessment, we compared wild type females mated to wild type males with wild type females mated to *iab-6^cocu^*mutants (**Figure 1A**).

**Figure 1.**
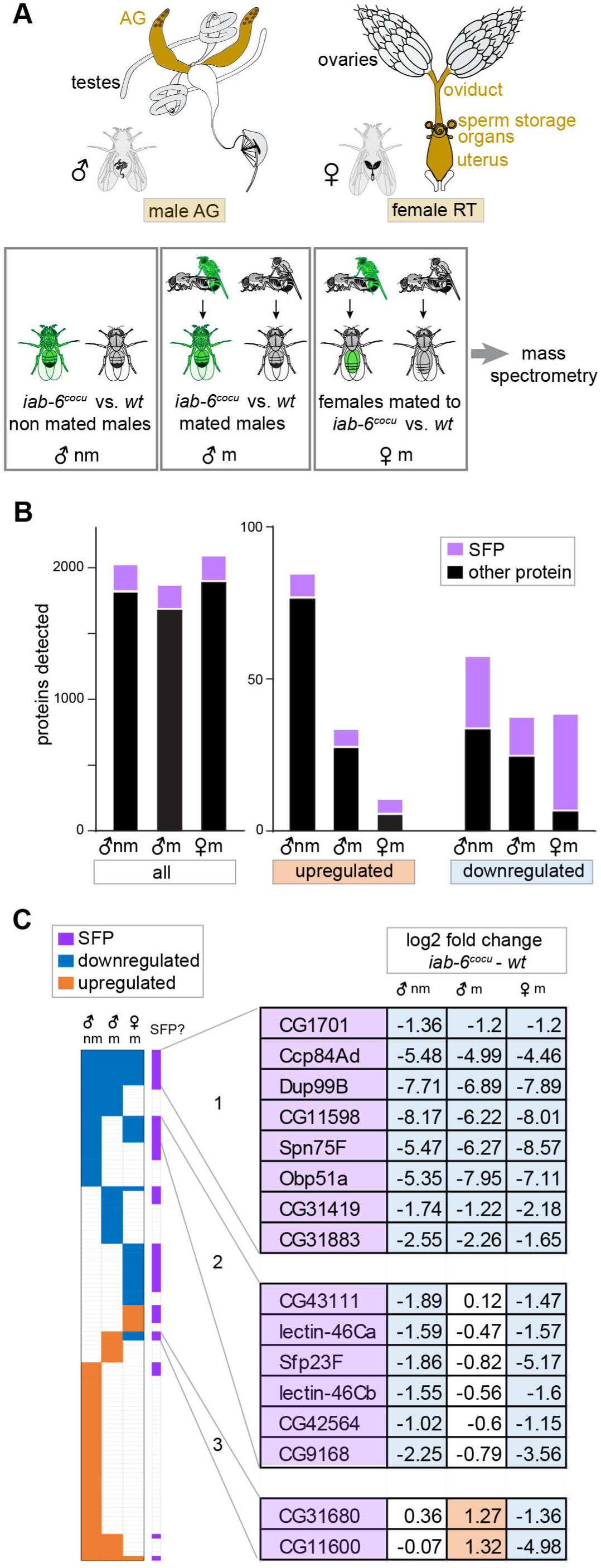
Downregulation of the seminal fluid transferome in *iab-6^cocu^* mutants. **A** Experimental design of differential mass spectrometry of male and female reproductive tissues for determination of *iab-6^cocu^* mutant proteome and transferome changes **B** Number of proteins detected, upregulated and downregulated in *iab-6^cocu^* mutants/mutant mates vs wild type controls. The three datasets are AG proteome non mated males (♂nm), AG proteome mated males (♂m) and non mated male, and RT proteome mated female (♀m). SFPs are shown in lilac. **C** List of high confidence transferome changes, i.e. proteins downregulated in the RT proteome mated female (♀m) and consistently changed in male proteomes (♂nm and ♂m), ordered based on hierarchical clustering into three groups. Numbers give the log2 fold change of protein abundance in *iab-6^cocu^* - wild type samples. Instances of log2 fold change≤-1 and adjusted p-value < 0.05 are indicated in blue (significantly downregulated in *iab-6^cocu^* samples), instances of log2 fold change ≥1 and adjusted p-value < 0.05 (significantly upregulated in *iab-6^cocu^* samples) are indicated in orange. SFPs are shown in lilac. For complete list, see Supplementary Table 2.

Across the three datasets, we identified 2037, 1882, and 2104 proteins with high confidence at the peptide and protein levels (1% FDR). Of the identified proteins, 85, 34 and 11 are significantly more abundant in *iab-6^cocu^*samples and 58, 38 and 39 are significantly less abundant (adjusted p-value < 0.05, 2 ≥ fold change) (**Figure 1B**, **Supplementary Table 1**).

In the three sample pairs, we identified 209, 188 and 199 of a total of 292 known high-confidence SFPs (Wigby et al. 2020). While known SFPs amount to approximately 10% of the total number of identified proteins, they are significantly enriched among the differentially abundant SFPs, constituting 22% (n = 32), 26% (n=19) and 74% (n=37) (Odds ratios of 3.08, 3.48 and 33.2, p < 0.0001, Fisher’s exact test) of the differentially abundant proteins in the AG proteome of mated and unmated males and in the mated female RT proteome, respectively (**Figure 1B**, **Supplementary Table 1**).

Differentially abundant proteins in the mated female RT could be due either to transferome changes in *iab-6^cocu^* mutant males or to differential translation/ degradation of female proteins in response to receiving a mutant *iab-6^cocu^*transferome. To address transferome changes, we considered proteins detected in all three datasets (n=1158) and focused on those that showed significant abundance changes in the mated female RT proteome and in one or both male proteomes, consistent with a transferred SFP. A single protein (Acp26Ab) is increased in all three datasets. Among proteins significantly decreased in the mated female RT proteome, we considered three groups. Group 1 proteins are significantly decreased in both male proteomes (n = 8). Group 2 proteins are decreased in the unmated male proteome and unchanged in the mated male proteome (n = 6). Group 3 proteins are unchanged in the unmated male proteome and increased in the mated male proteome (n=2). All 17 proteins that meet these criteria are high-confidence SFPs as classified by Wigby et al. (2020) (**Fig. 1C**, **Supplementary Table 2**).

We conclude that group 1 and 2 proteins are found in lower abundances in females mated to *iab-6^cocu^* males because they are present in lower abundances in AGs of *iab-6^cocu^* males. Group 3 proteins are most likely not affected in their abundances in *iab-6^cocu^* males, but rather in their ability to be transferred to the female. We assume that SFPs from groups 1-3 represent direct male transferome changes rather than a differential female response.

### SFPs depleted in the *iab-6 ^cocu^* mutant transferome affect female post-mating receptivity

Next, we wondered if SFPs from group 1-3 that were downregulated in the transferome were implicated in phenotypes associated with *iab-6^cocu^* males, namely their incapacity to trigger acoustic behavior in female mates during copulation (Kerwin et al. 2020) and failure to suppress female receptivity to remating (Gligorov et al. 2013).

We tested 15 of the 16 downregulated SFPs for which RNAi lines were available at public stock centers, using a ubiquitous driver line (*tubulin-GAL4*) for knockdown of gene expression in male flies and recorded female copulation song as well as remating propensity of their wild type mates. Depletion of the candidate SFPs by RNAi does not cause a significant decrease in female copulation song in mates of experimentally manipulated males for any of the lines tested (**Supplementary Figure 1**).

Among the downregulated SFPs, Lectin-46Ca, Lectin-46Cb and CG42564 have been previously reported to be required for suppression of female receptivity to remating (Ravi Ram and Wolfner 2007, Singh et al. 2018, Carlisle et al.2025). Among the remaining candidates, we find that ubiquitous knockdown of *CG1701* leads to a large proportion of females remating at 4d after the first mating when tested with a single wild type male for 1h (76%, n=44/58 and 65%, n=33/51, with two independent RNAi lines, respectively as compared to 0%, n=0/30 for the genetic driver line control).

We next assessed downregulated proteins in the mated female RT proteome, which did not show consistent changes in the male proteomes (n=12) or were not detected in both male proteomes (n=4) but were high-confidence SFPs (**Supplementary Table 1**). Recent studies have shown that around 40% of SFP transcripts are also expressed in the female RT (Thayer et al. 2024, Cridland and Begun 2023, McDonough-Goldstein et al. 2021). Three of the 16 proteins downregulated in the mated female RT proteome (CG18135, Cpr67Fb and Spn42Dd) belong to these reproductive fluid proteins with shared expression in male and females and their downregulation could be due to a differential response of the female RT to *iab-6^cocu^* mutant mates. The rest, i.e. 13 proteins, are more likely to be part of the transferome. Among these proteins, two additional mediators of the post-mating receptivity response have been reported: Aquarius (Findlay et al. 2014) and CG9997 (Ravi Ram and Wolfner 2007). As Lectin-46Ca and Lectin-46Cb, Aquarius and CG9997 belong to a small group of so-called sex peptide network proteins that mediate binding of sex peptide to sperm, movement of sex peptide into female storage organs and sustained, sex peptide dependent post-mating responses in females (Ravi Ram and Wolfner 2009, Singh et al. 2018). The involvement of CG42564 in sex peptide sperm binding has not been experimentally tested. Still, the protein has been suggested to be a sex peptide network member based on evolutionary analysis of correlated gene copy variation (Carlisle et al. 2025).

We conclude that downregulation of four characterized sex peptide network proteins, as well as two new receptivity affecting proteins (CG1701and CG42564) in the transferome of *iab-6^cocu^* mutant males, is likely to contribute to increased post-mating receptivity of their female mates.

### Functional characterization of two new mediators of the long-term female post-mating response, *cornutus* (CG1701) and *hanrej (*CG42564)

CG1701 and CG42564 were found to be transferred from male to female in proteomics studies using isotope-labeled females and thus were identified as SFPs (Findlay et al. 2008, Findlay et al. 2009) but have not been functionally studied in detail so far.

Given the strong effects on female post-mating receptivity and the phenotypic similarity of their knockdowns to the *cocu* mutant allele of *iab-6*, we propose new names for the previously unnamed genes: *cornutus* (CG1701) and *hanrej (*CG42564). Single-cell transcriptomics data indicate that expression of both genes is enriched in the AG and predominates in main cells (Li et al. 2022, Majane et al. 2021). To explore cell-type specific requirements, we used GAL4 driver lines labeling both main and secondary cells (*dve-GAL4*, Minami et al. 2012), main cells exclusively (*antares-GAL4*, a CRIMIC Trojan-Gal4 line from a gene-specific library, Lee et al. 2018) and secondary cells exclusively (*dsx-Gal4*, Rideout et al. 2010) (**Figure 2A, Supplementary Figure 2A**) to knock down *cornutus* and *hanrej*. Wild type levels of *cornutus* and *hanrej* mRNA are required in secondary cells and main cells of the AG, respectively, to prevent remating of females mated to experimental males at 4d after the first mating (**Figure 2A**). To confirm specificity of the phenotype to downregulation of protein in the AG, we combined the *dve-GAL4* driver, which also expresses in the innervation of the male RT, with a pan-neuronal *nsyb-GAL80*, suppressing nervous system expression and obtained the same results for knockdown of *cornutus* (**Supplementary Figure 2B, C**). Concomitant with the increase of post-mating receptivity, there is a drastic drop in fecundity, measured by the number of eggs laid per female, at 4d after mating, in mates of *cornutus* and *hanrej* knockdown males (**Figure 2B**). At 4d after mating, mates of knockdown males have the same number of sperm in their storage organs (seminal receptacles and spermatheca) as mates of control males (**Figure 2C**). In the ovaries of mates of knockdown males, an accumulation of late-stage egg chambers is seen at 4d, but not 2d after mating (**Figure 2D**). We conclude that Cornutus and Hanrej are required for ovulation of mature eggs in the presence of stored sperm.

**Figure 2.**
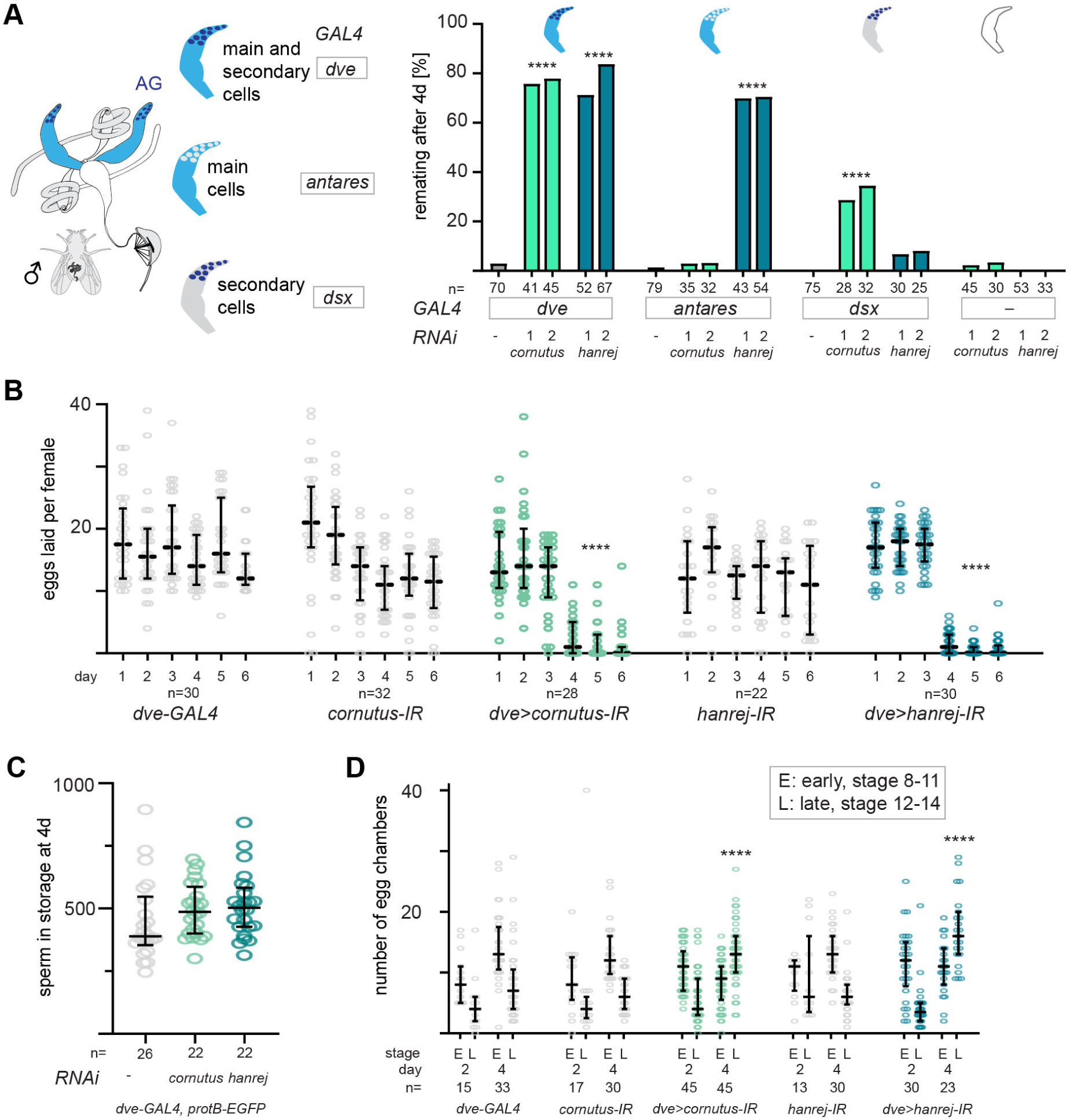
Cornutus and Hanrej are required for the sustained female post-mating response. **A** Schematic of driver lines used for expressing RNAi in different cell types of the AG (left). Percentage of remating female flies after 4d when mated to males in which *cornutus* o*r hanrej* were depleted with RNAi expressed in different AG cell populations and to control males (right). ****p < 0.0001, Fisher’s exact test. n, number of females tested, is indicated under the x axis. **B** Number of eggs laid by females mated to males in which *cornutus* o*r hanrej* were depleted and to control males, over the course of six days following the mating. ****p < 0.0001, Kruskal Wallis test with Dunn’s multiple comparison. **C** Number of sperm detected in female storage organs at 4d after mating for females mated to males in which *cornutus* o*r hanrej* were depleted with RNAi and to control males. No significant difference was detected between knockdown mates and the control (Mann-Whitney test, two sided). **D** Number of egg chambers, divided by E, early (8-11) and L, late (12-14) stages at day 2 and day 4 after mating found in females mated to males in which *cornutus* o*r hanrej* were depleted with RNAi and to control males. ****p < 0.0001, Kruskal Wallis test with Dunn’s multiple comparison. For full genotypes, see methods. Error bars represent median and interquartile range. n, number of females tested, is indicated under the x axis.

### Cornutus and Hanrej mediate binding of Sex peptide to sperm

The phenotypes observed upon *cornutus* and *hanrej* knockdown resemble closely the ones previously found for members of the sex peptide network (Singh et al. 2018). To test if Cornutus and Hanrej are required for the long-term association of SP with sperm, we assessed the presence of SP on sperm in the seminal receptacle by immunohistochemistry in females that were mated to males depleted for these proteins. At 4d after mating, the time point when females mated to knockdown males almost completely stop laying eggs, there is no detectable SP bound to sperm in these females, whereas females mated to control males have sperm with SP bound to its tails (**Figure 3**). Shortly after mating, at 6h after the end of copulation, when sperm storage is complete, sperm in seminal receptacles from males in which Cornutus or Hanrej were depleted show markedly weaker SP staining than sperm from control males (**Figure 3**). This leads us to conclude that Cornutus and Hanrej are required for efficient binding of SP to sperm tails shortly after mating and for the long-term presence of SP in the female receptacle.

**Figure 3.**
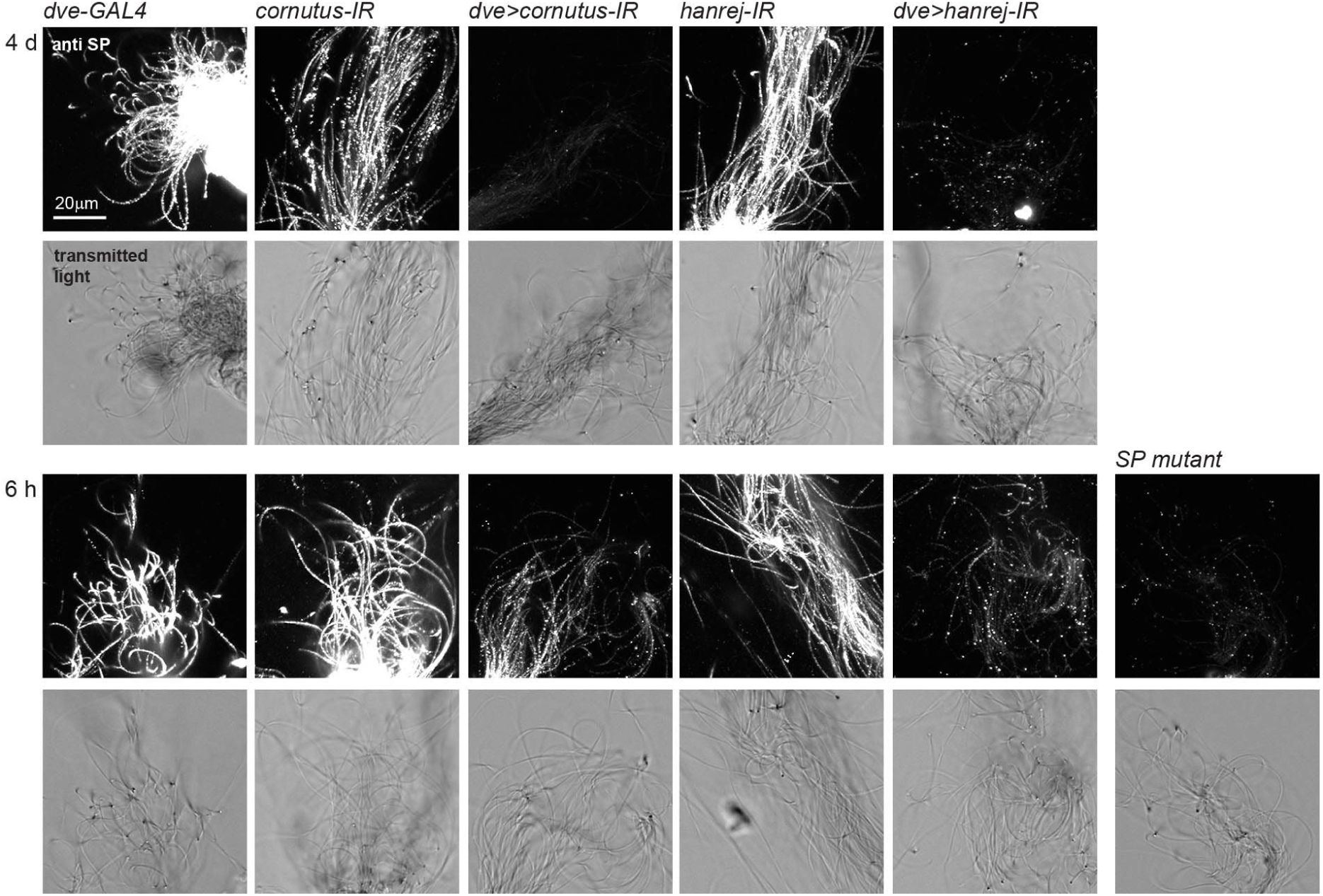
Cornutus and Hanrej are required for Sex peptide binding to sperm. Immunohistochemistry of SP (anti SP) on sperm dissected from seminal receptacles and visualized in transmitted light 4d (above) and 6h (below) after the female’s mating to males in which *cornutus* o*r hanrej* were depleted with RNAi and to control males, as well as sperm from a female mated to a *SP* mutant male. For full genotypes, see methods. Scale bar, 20μm. A micrograph of one representative sperm bundle (out of 3-8 samples per condition) is shown.

### Cornutus affects dense core granule maturation and secretion in accessory gland secondary cells

In *iab-6^cocu^* enhancer mutant males, the secondary cells of the AG lack large secretory vesicles (also called vacuoles) that are characteristic of this cell type (Gligorov et al. 2013). We used a newly described marker for secretory vesicles, mfas-Egfp, that accumulates in DCGs (Singh et al. 2025) to assess secondary cell morphology in knockdown males. Upon knockdown of either *cornutus* or *hanrej* in the accessory gland, we still observe mfas-Egfp positive DCGs within the large secretory vesicles of secondary cells (**Figure 4A**). However, DCG are smaller when Cornutus is depleted (**Figure 4B**). Together with the decreased size of DCGs in *cornutus* knockdown males, the mean number of DCGs increases per cell (p=0.001, Kruskal Wallis test with Dunn’s multiple comparison, comparing the mean DCG number of non-mated males, **Figure 4C**). Previously, it has been shown that a small number of mature DCGs are secreted from each secondary cell upon mating (Redhai et al. 2016). In agreement with this, we see around two mfas-Egfp labelled DCGs per cell being lost during mating in control flies. Upon knockdown of *cornutus*, the mean number of DCGs per cell did not differ before and after mating (**Figure 4C**). Such a loss of mating dependent DCG number reduction was not seen after knockdown of *hanrej* (data not shown). To assess the intracellular distribution of Cornutus, we expressed C-terminally Egfp-tagged Cornutus with the *dve-GAL4* driver. The tagged protein is detected in a subset of DCGs of secondary cells (**Figure 4D**). In contrast to mfas-Egfp, Cornutus-Egfp does not label DCGs in all secondary cells. Most Cornutus-Egfp positive DCG are present in very young males at the day of eclosion, i.e. a time point when the gland lumen has not yet fully filled with seminal fluid (**Figure 4D**). This indicates that Egfp-tagged Cornutus might accumulate in DCGs at a specific maturation state and later be processed or degraded. In summary, these results provide evidence that Cornutus is a protein required for proper DCG maturation and secretion in accessory gland secondary cells.

**Figure 4.**
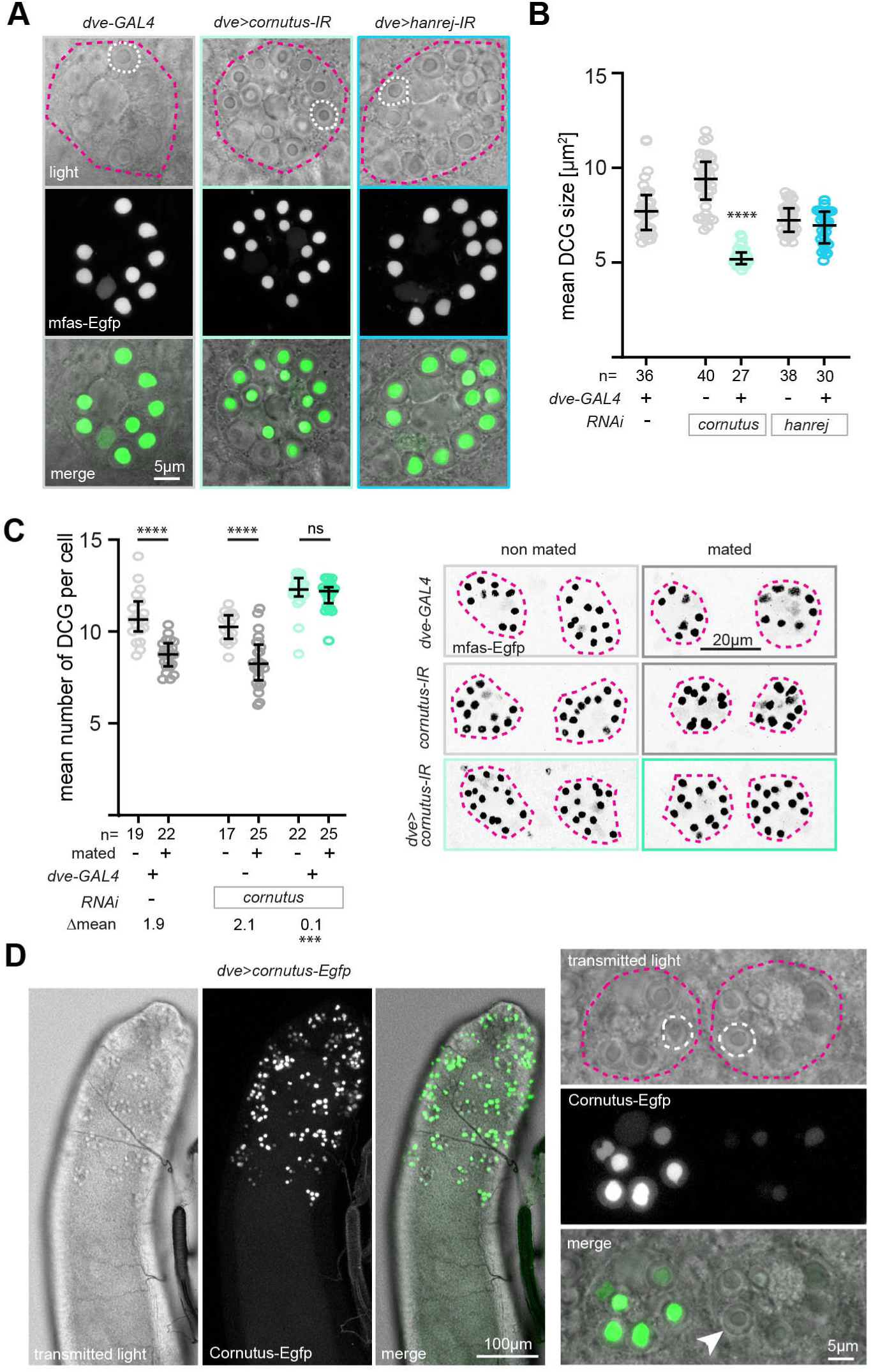
Cornutus is required for dense core granule maturation and release. **A** Representative images of secondary cells of control and knockdown males with mfas-Egfp labelled DCGs. In the transmitted light images, the circumference of the secondary cell is indicated by a pink dashed line, and one of the large secretory vesicles is outlined by a white dashed line. Scale bar, 5μm. **B** Mean size of mfas-Egfp labelled DCGs in control and knockdown males. n, number of secondary cells evaluated, is indicated under the x axis. ****p < 0.0001, Kruskal Wallis test with Dunn’s multiple comparison. **C** Mean number of DCGs per secondary cell in in control and knockdown males, for unmated and mated males. n, number of accessory glands evaluated, is indicated under the x axis. ****p < 0.0001, Mann Whitney test. The difference between unmated and mated males (Δmean) is indicated for each genotype. ***p=0.0006, permutation testing. On the right, DCGs of two representative secondary cells (mfas-Egfp in black, outline of cells indicated by pink dashed lines) are shown for each genotype and mating state. Scale bar, 20μm. For full genotypes, see methods. Error bars in **B, C** represent median and interquartile range. **D** Localization of Cornutus-Egfp, expressed with the *dve-GAL4* driver, in secondary cells DCGs. On the left, the distal part of an accessory gland is shown. On the right, adjacent two secondary cells (each indicated by a dashed pink line, with an exemplary large secretory vesicle outlined by a dashed white line) are shown, one with Cornutus-Egfp positive DCGs and one with faint or no Cornutus-Egfp label (white arrowhead) in DCGs. Scale bars, 100μm and 5μm.

## Discussion

### Abdominal-B expression in male accessory gland secondary cells shapes the seminal fluid transferome and transfer of Sex peptide network proteins

Female RT proteome analysis reveals that females mated to *iab-6^cocu^*mutants contain lower abundances of four well characterized SP network proteins, Lectin-46Ca, Lectin-46Cb, Aquarius and CG9997. Since each SP network protein is required individually for the female long-term response, combined downregulation of these four network proteins could be sufficient to explain the *iab-6^cocu^*mutant post-mating phenotype.

Overall, more than 10% of the known SFPs we detected in the AG and female RT proteome were differentially abundant in *iab-6^cocu^* mutants or females mated to these mutants, respectively. To stringently filter high confidence transferome changes in *iab-6^cocu^*mutants, we focused on proteins that were consistently changed in the data sets from unmated and/or mated males as well as in mated females, obtaining a list of 16 proteins with lower abundances in the RT of females mated to mutants, all of which were SFPs. Functional testing of these transferome-downregulated SFPs revealed two new SP network proteins, CG1701/Cornutus and CG42564/Hanrej, further strengthening the indication that SP network proteins are overrepresented among the SFPs downregulated in the *iab-6^cocu^*transferome.

Since none of these SP network proteins is downregulated in *iab-6^cocu^*mutants at the level of mRNA (Sitnik et al. 2016), it appears unlikely that this correlated depletion solely arises from transcriptional control. Rather, SP network members might depend on a shared mechanism for secretion, or post-secretion stabilization and transport that is orchestrated by Abd-B target genes in secondary cells.

There are indications that most *iab-6^cocu^* transferome changes stem from indirect mechanisms and that secondary cells, via their unique transcriptional and secretory profile, also modulate main cell SFP abundances and transfer. For example, we find two SFPs with a main cell expression bias (CG31680 and CG11600) significantly depleted in the female, present at normal levels in unmated mutant males, and elevated in mated mutant males, a pattern that clearly suggests failed transport. Only a single protein from our downregulated transferome list, CG11598, was previously reported to be strongly transcriptionally downregulated in *iab-6^cocu^* mutants (Sitnik et al. 2016). Single cell transcriptomics (Majane et al. 2022) suggests that among the 16 downregulated transferome SFP genes, only two, *lectin-46Ca* and *lectin-46Cb*, exhibit a strong expression bias in secondary cells. On the other hand, four genes are significantly enriched in expression in main cells and two in the anterior ejaculatory duct.

Taken together, the *iab-6^cocu^*transferome illustrates strong post-transcriptional control of SFP networks and interdependence of main and secondary cell products.

### Hanrej and Cornutus: New members of the Sex peptide network ensuring the long-term female post-mating response

Among the SFPs downregulated in the *iab-6^cocu^* mutant transferome, we find two new regulators of female fecundity and receptivity to remating, CG1701/Cornutus and CG42564/Hanrej. RNAi mediated knockdown of these two factors fully recapitulates the failure of the long-term post-mating response seen for *iab-6^cocu^*mutants or single characterized members of the SP network. In females mated to *cornutus* or *hanrej* knockdown males, production of fertilized eggs is normal for three days, but then drops sharply, coinciding with high remating rates at four days after mating. Females normally remate and stop laying eggs when they have depleted stored sperm (Manning 1967). When females mated to *cornutus* or *hanrej* knockdown males loose post-mating behavior when they still have a higher number of sperm in their storage organs. However, they do not utilize this stored sperm for fertilization and accumulate mature oocytes in their ovaries. Such inability to detect and use sperm is a hallmark of defects in the SP network that mediates the binding of SP to sperm in the female RT (Avila et al. 2010, Singh et al. 2018). In mates of *cornutus* or *hanrej* knockdown males, SP on sperm tails is not detectable at 4d after mating and is already markedly reduced 6h after mating when sperm storage is completed. This indicates that Cornutus and Hanrej are required for the transfer of SP onto sperm tails, either by directly interacting with sperm themselves or by involvement in preceding SP network signaling steps.

We establish that *cornutus*, though judged by single cell transcriptional analysis as a main cell marker gene with strong expression bias (Majane et al. 2022), is not required in this cell type for the female long-term post-mating response, but rather in secondary cells, whereas *hanrej* is required in main cells. For other members of the SP network that transiently bind to sperm together with SP, an origin from secondary cells (Lectin-46Ca, Lectin-46Cb) as well as from main cells (Antares, CG9997) has been implied before. The SP network thus appears to rely on a tight cooperation of secondary and main cell secretory products.

### Secondary cell DCG vesicle secretion as a key step for producing a functional SP network

Unusually large DCG vesicles are characteristic of secondary cells. Though surpassing the size of typical mammalian DCG vesicles in the nervous system and hormone producing glands by more than an order of magnitude, they share evolutionary conserved regulators of biogenesis and trafficking (Wells et al. 2023, Singh et al. 2025). The large DGC vesicles not only contain and release material from DCGs, but also small intraluminal vesicles/exosomes (Corrigan et al. 2014) and electron dense filamentous structures (Bairati 1968, Rylett et al. 2007). Mating induced DCG vesicle release stimulates their replenishment via an autocrine mechanism involving BMP signaling (Redhai et al. 2016), and interference with this pathway has been previously shown to affect some aspects of female post-mating behavior and sperm handling, as well as the AG proteome (Hopkins et al. 2019). In secondary cells of *iab-6^cocu^* mutants, DCG vesicles are completely lost, along with other morphological changes such as cell shape and volume.

We find that loss of Cornutus specifically impairs the release of DCG vesicles during mating. In *cornutus* knockdown secondary cells, DCG vesicles still form, but are slightly more numerous and contain smaller condensed DCG cores in unmated males. This suggests that the final DCG maturation steps, aggregation of smaller minicores to a big central DCG within a vesicle (Singh et al. 2025) and triggered secretion of vesicles during copula require Cornutus, which itself localizes to DCG vesicles. Without Cornutus, the SP network does not function normally, and all canonical aspects of the long-term post-mating response are lost. Currently, it is not known if this SP network failure results from a lack of Cornutus transfer to the female RT or from the lack of other SP network proteins that are released by DGC vesicles as part of the DCGs, exosomes or filamentous structures. The main candidates for DCG vesicle release dependent SP network proteins are Lectin-46Ca and Lectin-46Cb.

More detailed studies of the interactions of previously known SP network members with the two newly identified factors, Cornutus and Hanrej, are needed to clarify which parts of the network depend directly on DCG vesicle secretion.

In conclusion, disrupted secondary cell secretory identity in *iab-6^cocu^*mutant males influences the abundances of more than 10% of known SFP in the AG. The vast majority of the differentially abundant SFPs are not under direct transcriptional control of Abd-B, but likely to be subjected to altered translation, processing and secretion, stability and/or transfer. In the future, it will be of interest to investigate the mechanisms by which secondary and main cells interact and the interdependence of their secretory products in the seminal fluid in both male and female RT. Such interdependence is of functional relevance within the SP network, which relies on SFPs from main and secondary cells. In the *iab-6^cocu^* mutant transferome, multiple members of the SP network are downregulated. The mechanism of this concerted regulation remains to be explored, as does the exact role of DCG vesicle products in mediating the long-term post-mating response.

## Methods

### Drosophila culture and transgenic Drosophila stocks

Flies were maintained on a standard medium containing sugar, oatmeal, cornmeal, agar, and yeast, under 12h light:12h dark cycle at 25°C. All experimental flies were collected after eclosion and aged 4-7 days. Males were individually aged in a flat bottom 1.5 mL 96 well block filled with 0.5 mL food per well and covered with a PCR foil with air holes. Females were aged in groups of 10-15 in food vials. Fly stocks used in this study are listed in **Supplementary Table 3**. The exact genotypes of experimental flies are listed in **Supplementary Table 4**. To generate *UAS-cornutus-Egfp* flies, the protein coding sequence with a C-terminal Egfp tag of cornutus was cloned into the *pUASTattB* vector and the construct was inserted by PhiC31 mediated integration into the *attp40* landing site.

### Mass Spectrometry

A male (for unmated sample) or a male and a virgin female (for mated samples) were transferred to a perforated 1.5 mL Eppendorf tube and kept under observation until the occurrence of copulation. Immediately after the end of copulation (for the mated group), flies were frozen in liquid nitrogen. Tissues were dissected with fine forceps in ice cold PBS, transferred to a 1.5mL Eppendorf tube and frozen in liquid nitrogen. Each sample contained tissues from five individual flies, and each condition was analyzed in five biological replicates. Male tissues were the paired accessory glands, removed from the ejaculatory duct at their bases. Female tissues were the reproductive tract with the ovaries removed at the base of the lateral oviducts. Tissues were lyophilized dry before being resuspended in 8M urea. As urea was added, the tissues were thoroughly macerated with the tip of the pipette. Samples were reduced for 60min in 10mM dithiothreitol, followed by alkylation for 60min in 30mM iodoacetamide. The urea concentration was lowered to <1 M before overnight digestion with trypsin (2 μg/sample) at 37°C. The samples were desalted on homemade columns packed with octadecyl C18 disks (Empore, 3M) and dissolved in 0.1% formic acid. The desalted samples were loaded onto a trap column (2cm x 75μm inner diameter) and separated on an analytical column (15 cm x 75μm inner diameter) packed with 3 μm C18 beads (Dr. Maisch, GmbH). Samples were eluted over 60min and analyzed on an Orbitrap Eclipse Tribid mass spectrometer (Thermo Fisher Scientific). Samples were analyzed in technical triplicates and injected in alternating order. Data files were processed in Proteome Discoverer 2.4 with the Sequest HT search engine. The *Drosophila melanogaster* UniProt reference proteome (UP000000803) was used as a database, with the following parameters: 10 ppm precursor mass accuracy, 0.6 Da fragment mass accuracy, trypsin as digestion enzyme, maximum three missed cleavage sites, Oxidation (M) as variable modification and Carbamidomethyl (C) as fixed modification. Unique peptides were quantified using a label-free approach based on precursor intensity or area to compare protein abundances across samples. Protein abundances were normalized to total peptide amount.

### Mass spectrometry data analysis

Data were filtered to only include proteins identified with a 1% FDR at the peptide level and to only include proteins identified in at least 4 out of 5 biological replicates (in either condition). The median of the technical replicates was used for further analyses. Protein abundances were base 2 log transformed before further processing. Missing values were imputed using a probabilistic minimum method. A Gaussian distribution was fit to the log2 transformed abundances, downshifted by 1.8 standard deviations and normalized to have a 30% standard deviation, to simulate the detection threshold of the mass spectrometer. Missing values were then drawn at random from this distribution. The imputation was performed 1000 times, and only proteins significantly different in abundance in 60% of the imputations were used to specify differently abundant proteins. P-values were calculated using an unpaired equal variance two-sample t-test and FDR-corrected using the Benjamini-Hochberg procedure (Benjamini and Hochberg, 1995). Fold change was calculated as the mean of the pairwise fold changes between log2 abundances of *iab-6^cocu^* and wild type samples. Proteins were considered differentially abundant at an absolute log2 fold change of ≥1 and an adjusted p-value< 0.05. Hierarchical clustering was performed using the MATLAB function clustergram, with average linkage and Euclidean distances on standardized protein abundances.

### Behavioral assays

Behavioral assays were conducted at 25°C, 60% humidity at circadian time 0-3hrs after lights on or 3-0hrs before lights off. Female copulation song was recorded and analyzed as previously described (von Philipsborn et al. 2023a). Mating assays and tests of post-mating receptivity were performed as previously described (von Philipsborn et al. 2023b, c). Female virgin flies were paired with experimental males for 1h in circular mating chambers with 10mm diameter, 4mm height. If mating occurred, females were collected and kept in groups of 5-10 flies in food vials for 4 days. Remating was tested with wild type males for 1h in mating chambers. For assessing egg laying, virgin females were individually mated in mating chambers and then individually housed in 5ml vials filled with 1ml of fly food. Every 24hrs, females were transferred to new vials and eggs laid were counted.

### Sperm and egg chamber counts

To quantify female sperm storage, the spermatheca and seminal receptacle of mated females were dissected in 4°C PBS and mounted in a drop of PBS on a glass slide by pressing down gently a coverslip onto the organs. ProtamineB-Egfp labeled sperm heads were imaged with a fluorescent microscope (Nikon Eclipse Ni, with 20x/0.5 and 40x/0.75 Plan Fluor Air Objectives) and counted using the Fiji/ImageJ cell counter plug-in. To quantify egg chambers, ovaries of mated females were dissected in 4°C PBS, fixed in 4% paraformaldehyde with 0.3% TritonX for 25min at room temperature, washed three times in PBS with 0.3% TritonX and mounted in Vectashield containing Dapi (Vector labs, RRID: AB_2336790). Ovaries were imaged with a Leica STELLARIS8 FALCON confocal microscope equipped with a HC PL APO 20x/0,75 multi-immersion objective. Egg chambers were staged based on nuclear staining (Jia et al. 2016).

### Immunohistochemistry

Immunohistochemistry of fly nervous systems and reproductive tracts was performed as described previously (Amin et al. 2023). Tissues were dissected in 4°C PBS, fixed in 4% paraformaldehyde with 0.3% TritonX, stained with mouse nc82 antibody (Developmental Studies Hybridoma Bank, RRID:AB_2314866, for nervous systems), rabbit anti-GFP antibody (Torrey Pines Biolabs, RRID: AB_10013661) followed by respective secondary antibodies (Goat IgG (H+L) Alexa Fluor 488 and 647) and Alexa Fluor 647 conjugated Phalloidin (Thermo Fisher, RRID: AB_2620155, for reproductive tracts) and mounted in Vectashield (Vector labs, RRID: AB_2336789).

Immunohistochemistry of sperm-bound SP was performed as described previously (Misra and Wolfner 2020). Seminal receptacles of mated females were dissected on poly-L-lysine coated glass slides 4°C PBS, fixed in 4% paraformaldehyde for 15min at room temperature, washed three times in PBS, blocked for for 30 minutes in 5% normal goat serum (NGS) and then incubated overnight at 4°C with rabbit anti-SP primary antibody (M. Wolfner lab, 1:200), followed by incubation for 2-3hrs at room temperature with secondary antibody (Goat IgG (H+L) Alexa Fluor 488). Samples were mounted in Vectashield (Vector labs, RRID: AB_2336789). Dissection and staining steps were carried out within a ring of double-sided adhesive tape attached to the glass slide to confine the sample area and prevent sample loss. Tissues and sperm bundles were imaged with a Leica STELLARIS8 FALCON microscope equipped with a HC PL APO 20x/0,75 multi-immersion objective.

### DCG imaging data analysis

For imaging of mfas-Egfp and Cornutus-Egfp, accessory glands were dissected in 4°C PBS and mounted between two coverslips in a drop of PBS, with a small amount of food-grade silicone grease serving as spacer between the coverslips. The coverslips were inserted in a metal frame and imaged with a Leica STELLARIS8 FALCON microscope (HC PL APO 100x/1,40 oil objective for assessing DCG size, HC PL APO 20x/0,75 multi-immersion objective for counting DCG). To assess the size of the DCG we used the plug-in “analyse particle” in FIJI software. For each genotype 4 animals with 3-6 secondary cells per accessory gland were evaluated.

To assess DCG number, they were counted based on manual inspection of z-stacks using merged fluorescence and transmitted light channels. Approximately 10 secondary cells per AG were analysed. Only condensed DCGs were included in the analysis, characterized by sharply defined circular structures with a bright GFP signal. Mated flies were anesthetized on ice immediately after the end of copulation and their accessory glands were imaged within 2hrs. For permutation testing of DGC release, the differences between the mean number of DCG per cell (Δmean) before and after mating were calculated. All before mating datapoints for different genotypes and all after mating datapoints for different genotypes were permuted 100.000 times, the p-value reports the probability of the difference of the absolute Δmean for the knockdown genotype to the Δmeans of both control genotypes being ≥ the observed value (code deposited under: github.com/KinaUna/PhilipsbornPermutations).

## Data availability

The mass spectrometry proteomics data have been deposited to the ProteomeXchange Consortium (http://proteomecentral) via the PRIDE (Perez-Riverol et al. 2025) partner repository with the dataset identifier PXD072877.

## Acknowledgements

We thank Anna Prudnikova, Valerie Salicio, Laurence Clément, Isabelle Scerri and David Rodriguez Crespo for technical assistance. We thank DGRC and VDRC stock libraries, Rob Maeda, Jean Christophe Billeter and Hideki Nakagoshi for providing fly stocks, Developmental Studies Hybridoma Bank for the nc82 antibody, Mariana Wolfner for the anti-SP antibody. This study was supported by Lundbeckfonden grant DANDRITE-R248-2016-2518 and by Swiss National Science foundation grant 310030_212222.

## Supplementary Data

### Supplementary Figures

**Supplementary Figure 1.**
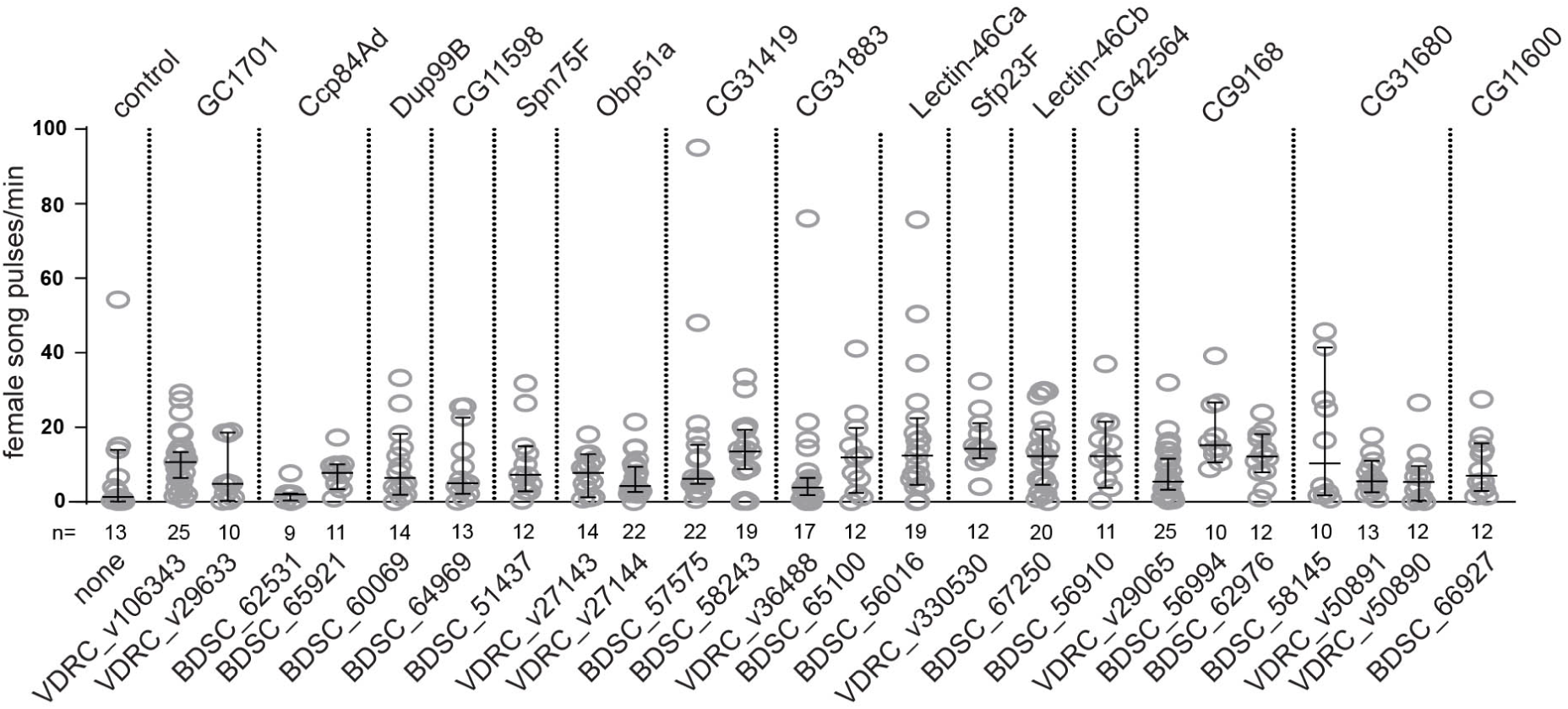
Copulation song of females mated to males depleted for candidate SFPs downregulated in the *iab-6^cocu^* mutant transferome. Female copulation song pulses/ min copulation of wild type females mated to experimental males, in which candidate SFPs were depleted by RNAi expressed by the ubiquitous *tubulin-GAL4* driver. SFP names are shown above the graph, identifiers of the RNAi lines under the graph. Each datapoint represents one copulating female, with n indicated under the x axis. Error bars represent median and interquartile range, no significant difference was detected to the control (Mann-Whitney test, two sided).

**Supplementary Figure 2.**
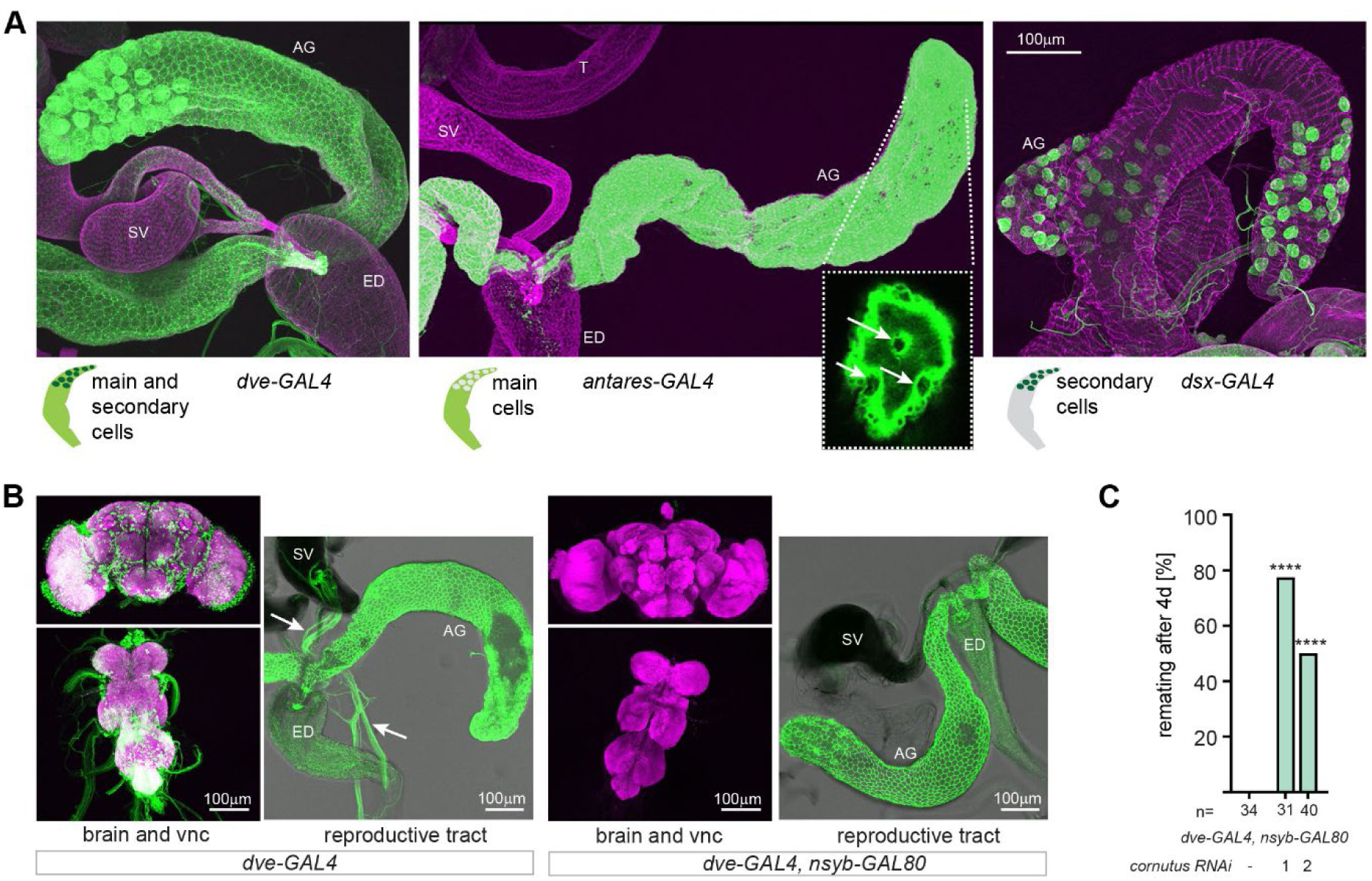
Expression patterns of GAL4 driver lines to specifically manipulate main and secondary cells of the accessory gland. A Expression pattern of *dve-GAL4*, *antares-GAL4* and *dsx-GAL4* in the male reproductive tract, cd8-Egfp in GAL4 expressing cells is shown in green, phalloidin based actin staining in muscle is shown in magenta. The inset in the middle panel shows a single confocal imaging plane with non-labelled secondary cells indicated by white arrows. B Expression pattern of dve-GAL4 without (left) and with (right) suppression of neuronal expression by *nsyb-GAL80* in central nervous system (brain and vnc) and reproductive tract. cd8-Egfp in GAL4 expressing neurons and gland cells is shown in green, anti-bruchpilot staining of the central nervous system neuropil in magenta, transmitted light outlines of the reproductive tract in greyscale. A micrograph of one representative tissue (out of 5 or more samples per genotype with the same expression pattern) is shown. AG, accessory gland, SV, seminal vesicle, ED, ejaculatory duct, T, testis. Scale bar, 100μm. C Percentage of remating female flies after 4d when mated to males in which *cornutus* was depleted with RNAi expressed by the *dve-GAL4, nsyb-GAL80* driver. ****p < 0.0001, Fisher’s exact test.

### Supplementary Tables

**Supplementary Table 1 (Excel file)**

The three sheets list detected proteins for 1) AG proteome non mated males (male nm), 2) AG proteome mated males (male m), and 3) RT proteome mated female (female m). Significantly up- or downregulated proteins in *iab-6^cocu^* mutants or females mated to *iab-6^cocu^* mutants are marked with “1” in column I and J, high-confidence SFPs are marked with “1” in column W. Columns K-O and P-T list protein abundancies of the 5 biological replicates (containing each tissues from 5 flies) for *iab-6^cocu^* mutants (cocu) and wild type samples (wt).

**Supplementary Table 2 (Excel file)**

List of proteins detected in all three datasets with three groups of proteins that show significant abundance changes consistent with a downregulation in the male transferome (see Figure 1C).

**Supplementary Table 3.** Drosophila stocks used.

| Short name, label | Genotype | Identifier/RRID |
| --- | --- | --- |
| wt, CS | Canton S wild type strain | Laboratory of B. Dickson |
| *iab-6<sup>cocu</sup>* | +; +; *iab-6<sup>cocu</sup>*<br>Mutant in wt background | Laboratory of F. Karch, R. Maeda, FBal0284736 |
| *tub-GAL4* | *y, w; P{tubP-GAL4}LL7* | BDSC_5138 |
| *dve-GAL4* | *w;;P{dve-GAL4.13}30A* | Laboratory of H. Nakagoshi, FBti0152837 |
| *antares-GAL4* | *y, w; Tl{CRIMIC.TG4.2}antr<sup>CR00860-TG4.2</sup>* | FBal0341147 |
| *dsx-GAL4* | *w;;Tl{GAL4}dsx[GAL4]/TM6B, Tb* | BDSC_66674 |
| *nsyb-GAL80* | *w; P{w[+mC]=nSyb-GAL80.S}3* | BDSC_92154 |
| *protB-eGFP* | +; *P{protamineB-eGFP}2* | Derived from BDSC_58406 |
| *SP mutant* | *SP0(0325)* | BDSC_94694 |
| *SP deficiency* | *Df(3L)Δ130* | Laboratory of J.C. Billeter, FBab0029476 |
| *UAS-Dcr2* | *w, P{UAS-dicer2, w[+]}* | VDRC Id 60007 |
| *RNAi: CG1701, cornutus-IR, 1* | *w; P{KK109689}VIE-260B* | VDRC Id 106343 |
| *RNAi: CG1701, cornutus-IR, 2* | *w; P{GD15054}v29663* | VDRC Id 29663 |
| *RNAi: CG42564, hanrej-IR, 1* | *y, sc, v, sev; P{TRiP.HMC06173}attP40* | BDSC_65910 |
| *RNAi: CG42564, hanrej-IR, 2* | *w;; P{GD6109}v14158* | VDRC Id 14158 |
| *mfas-Egfp* | *y, w; Mi{PT-GFSTF.2}mfas[MI11275-GFSTF.2]* | BDSC_63204 |
| *UAS-cornutus-Egfp* | *w; P{UAS-cornutus-Egfp}attP40* | This study |
| *UAS-cd8-Egfp* | *w; P{UAS-cd8-Egfp}* | Laboratory of B. Dickson |
| *RNAi: Ccp84Ad-IR, 1* | *y, v; P{TRiP.HMJ24075}attP40/CyO* | BDSC_62531 |
| *RNAi: Ccp84Ad-IR, 2* | *y, sc, v, sev; P{TRiP.HMC06176}attP40* | BDSC_65912 |
| *RNAi: Dup99B-IR* | *y, sc, v, sev; P{TRiP.HMC05062}attP40* | BDSC_60069 |
| *RNAi: CG11598-IR* | *y, sc, v, sev; P{TRiP.HMC05843}attP40* | BDSC_64969 |
| *RNAi: Spn75F-IR* | *y, v; P{TRiP.HMC03151}attP2* | BDSC_51437 |
| *RNAi: Obp51a-IR, 1* | *w; P{GD6816}v27143* | VDRC Id 27143 |
| *RNAi: Obp51a-IR, 2* | *w; P{GD6816}v27144* | VDRC Id 27144 |
| *RNAi: CG31419-IR, 1* | *y, sc, v, sev; P{TRiP.HMC04584}attP40* | BDSC_57575 |
| *RNAi: CG31419-IR, 2* | *y, v; P{TRiP.HMJ22264}attP40/CyO* | BDSC_58243 |
| *RNAi: CG31883-IR, 1* | *y, sc, v, sev; P{TRiP.HMC05996}attP40* | BDSC_65100 |
| *RNAi: CG31883-IR, 2* | *w; P{GD14757}v36488* | VDRC Id 36488 |
| *RNAi: lectin-46Ca-IR* | *y, sc, v, sev; P{TRiP.HMC04312}attP40* | BDSC_56016 |
| *RNAi: Sfp23F-IR* | *w; P{VSH330530}attP40* | VDRC Id 330530 |
| *RNAi: lectin-46Cb-IR* | *y, sc, v, sev; P{TRiP.HMC06352}attP40* | BDSC_67250 |
| *RNAi: CG9168-IR, 1* | *w; P{GD14142}v29065* | VDRC Id 29065 |
| *RNAi: CG9168-IR, 2* | *y, sc, v, sev; P{TRiP.HMC04436}attP40* | BDSC_56994 |
| *RNAi: CG9168-IR, 3* | *y, v; P{TRiP.HMJ30053}attP40* | BDSC_62976 |
| *RNAi: CG31680-IR, 1* | *y, v; P{TRiP.HMJ22095}attP40* | BDSC_58145 |
| *RNAi: CG31680-IR, 2* | *w; P{GD17288}v50891* | VDRC Id 50891 |
| *RNAi: CG31680-IR, 3* | *w; P{GD17288}v50890* | VDRC Id 50890 |
| *RNAi: CG11600-IR* | *y, sc, v, sev; P{TRiP.HMS05393}attP40* | BDSC_66927 |

**Supplementary Table 4.** Genotypes of experimental flies. All genotypes refer to male flies. All female mates used in experiments were CS wild type. Chromosomes derived from CS wild type background are written “*+*”.

| Figure | label | Full genotype |
| --- | --- | --- |
| 2A | *dve*-GAL4 | *w*, *UAS-Dcr2*; +/+; *dve*-GAL4/+ |
|  | *dve*-GAL4, *cornutus*-IR, 1 | *w*, *UAS-Dcr2*; P{KK109689}VIE-260B/+; *dve*-GAL4/+ |
|  | *dve*-GAL4, *cornutus*-IR, 2 | *w*, *UAS-Dcr2*; P{GD15054}v29663/+; *dve*-GAL4/+ |
|  | *dve*-GAL4, *hanrej*-IR, 1 | *w*; P{TRiP.HMC06173}attP40/+; *dve*-GAL4/+ |
|  | *dve*-GAL4, *hanrej*-IR, 2 | *w*, *UAS-Dcr2*; +/+; *dve*-GAL4/P{GD6109}v14158 |
|  | *antares*-GAL4 | *w*, *UAS-Dcr2*; *antares*-GAL4/+; +/+ |
|  | *antares*-GAL4, *cornutus*-IR, 1 | *w*, *UAS-Dcr2*; *antares*-GAL4/P{KK109689}VIE-260B; +/+ |
|  | *antares*-GAL4, *cornutus*-IR, 2 | *w*-, *UAS-Dcr2*; *antares*-GAL4/P{GD15054}v29663; +/+ |
|  | *antares*-GAL4, *hanrej*-IR, 1 | +; *antares*-GAL4/P{TRiP.HMC06173}attP40/+; +/+ |
|  | *antares*-GAL4, *hanrej*-IR, 2 | *w*, *UAS-Dcr2*; +/+; *dve*-GAL4/P{GD6109}v14158 |
|  | *dsx*-GAL4 | *w*, *UAS-Dcr2*; +/+; *dsx*-GAL4/+ |
|  | *dsx*-GAL4, *cornutus*-IR, 1 | *w*, *UAS-Dcr2*; P{KK109689}VIE-260B/+; *dsx*-GAL4/+ |
|  | *dsx*-GAL4, *cornutus*-IR, 2 | *w*, *UAS-Dcr2*; P{GD15054}v29663/+; *dsx*-GAL4/+ |
|  | *dsx*-GAL4, *hanrej*-IR, 1 | *w*; P{TRiP.HMC06173}attP40/+; *dsx*-GAL4/+ |
|  | *dsx*-GAL4, *hanrej*-IR, 2 | *w*, *UAS-Dcr2*; +/+; *dsx*-GAL4/P{GD6109}v14158 |
|  | *cornutus*-IR, 1 | +; P{KK109689}VIE-260B/+; +/+ |
|  | *cornutus*-IR, 2 | +; P{GD15054}v29663/+; +/+ |
|  | *hanrej*-IR, 1 | +; P{TRiP.HMC06173}attP40/+; +/+ |
|  | *hanrej*-IR, 2 | +; +/+; P{GD6109}v14158/+ |
| 2C, E | *dve*-GAL4 | *w*, *UAS-Dcr2*; +/+; *dve*-GAL4/+ |
|  | *cornutus*-IR | +; P{KK109689}VIE-260B/+; +/+ |
|  | *dve*>*cornutus*-IR | *w*, *UAS-Dcr2*; P{KK109689}VIE-260B/+; *dve*-GAL4/+ |
|  | *hanrej*-IR | +; +/+; P{GD6109}v14158/+ |
|  | *dve*>*hanrej*-IR | *w*, *UAS-Dcr2*; +/+; *dve*-GAL4/P{GD6109}v14158 |
| 2D | *dve*-GAL4, *protB*-Egfp | *w*, *UAS-Dcr2*; *protamineB*-Egfp/+; *dve*-GAL4/+ |
|  | *dve*-GAL4, *protB*-Egfp, *cornutus* RNAi | *w*, *UAS-Dcr2*; P{KK109689}VIE-260B/*protamineB*-Egfp; *dve*-GAL4/+ |
|  | *dve*-GAL4, *protB*-Egfp, *hanrej* RNAi | *w*, *UAS-Dcr2*; *protamineB*-Egfp/+; *dve*-GAL4/P{GD6109}v14158 |
| 3 | *dve*-GAL4 | *w*, *UAS-Dcr2*; +/+; *dve*-GAL4/+ |
|  | *cornutus*-IR | +; P{KK109689}VIE-260B/+; +/+ |
|  | *dve*>*cornutus*-IR | *w*, *UAS-Dcr2*; P{KK109689}VIE-260B/+; *dve*-GAL4/+ |
|  | *hanrej*-IR | +; P{TRiP.HMC06173}attP40/+; +/+ |
|  | *dve*>*hanrej*-IR | *w*, *UAS-Dcr2*; P{TRiP.HMC06173}attP40/+; *dve*-GAL4/+ |
|  | *SP* mutant | +; +/+; *SP0*(0325)/ *Df*(3L)Δ130 |
| 4A, B, C | *dve-GAL4* | *w, UAS-Dcr2; +/+; dve-GAL4/mfas-Egfp* |
|  | *cornutus-IR* | *+*; *P{KK109689}VIE-260B/+; +/+* |
|  | *dve>cornutus-IR* | *w, UAS-Dcr2; P{KK109689}VIE-260B/+; dve-GAL4/ mfas-Egfp* |
|  | *hanrej-IR* | *+*; *P{TRiP.HMC06173}attP40/+; +/+* |
|  | *dve>hanrej-IR* | *w, UAS-Dcr2; P{TRiP.HMC06173}attP40/+; dve-GAL4/+* |
| 4D | *dve>cornutus-Egfp* | *w; P{UAS-cornutus-Egfp}attP40/+; dve-GAL4, nsyb-GAL80/+* |
| Supp. 1 | control | *+*; *+/+; tub-GAL4/+* |
|  | CG1701, VDRC_106343 | *+*; *P{KK109689}VIE-260B/+; tub-GAL4/+* |
|  | CG1701, VDRC_29663 | *+*; *P{GD15054}v29663/+; tub-GAL4/+* |
|  | Ccp84Ad, BDSC_62531 | *+*; *P{TRiP.HMJ24075}attP40/+; tub-GAL4/+* |
|  | Ccp84Ad, BDSC_65912 | *+*; *P{TRiP.HMC06176}attP40/+; tub-GAL4/+* |
|  | Dup99B, BDSC_60069 | *+*; *P{TRiP.HMC05062}attP40/+; tub-GAL4/+* |
|  | CG11598, BDSC_64969 | *+*; *P{TRiP.HMC05843}attP40/+; tub-GAL4/+* |
|  | Spn75F, BDSC_51437 | *+*; *+/+; tub-GAL4/P{TRiP.HMC03151}attP2* |
|  | Obp51a, VDRC_27143 | *+*; *P{GD6816}v27143/+; tub-GAL4/+* |
|  | Obp51a, VDRC_27144 | *+*; *P{GD6816}v27144/+; tub-GAL4/+* |
|  | CG31419, BDSC_57575 | *+*; *P{TRiP.HMC04584}attP40/+; tub-GAL4/+* |
|  | CG31419, BDSC_58243 | *+*; *P{TRiP.HMJ22264}attP40/+; tub-GAL4/+* |
|  | CG31883, BDSC_65100 | *+*; *P{TRiP.HMC05996}attP40/+; tub-GAL4/+* |
|  | CG31883, VDRC_36488 | *+*; *+/+; P{GD14757}v36488/ tub-GAL4* |
|  | lectin-46Ca, BDSC_56016 | *+*; *P{TRiP.HMC04312}attP40/+; tub-GAL4/+* |
|  | Sfp23F, VDRC_330530 | *+*; *P{VSH330530}attP40/+; tub-GAL4/+* |
|  | lectin-46Cb, BDSC_67250 | *+*; *P{TRiP.HMC06352}attP40/+; tub-GAL4/+* |
|  | CG9168, VDRC_29065 | *+*; *P{GD14142}v29065/+; tub-GAL4/+* |
|  | CG42564, VDRC_14158 | *+*; *+/+; P{GD6109}v14158/ tub-GAL4* |
|  | CG9168, BDSC_56994 | *+*; *P{TRiP.HMC04436}attP40/+; tub-GAL4/+* |
|  | CG9168, BDSC_62976 | *+*; *P{TRiP.HMJ30053}attP40/+; tub-GAL4/+* |
|  | CG31680, BDSC_58145 | *+*; *P{TRiP.HMJ22095}attP40/+; tub-GAL4/+* |
|  | CG31680, VDRC_50891 | *+*; *+/+; P{GD17288}v50891/tub-GAL4* |
|  | CG31680, VDRC_50890 | *+*; *+/+; P{GD17288}v50890/tub-GAL4* |
|  | CG11600, BDSC_66927 | *+*; *P{TRiP.HMS05393}attP40/+; tub-GAL4/+* |
| Supp. 2A, B | *dve-GAL4* | *w; UAS-cd8-Egfp/+; dve-GAL4/+* |
|  | *antares-GAL4* | *w; UAS-cd8-Egfp/antares-GAL4; +/+* |
|  | *dsx-GAL4* | *w; UAS-cd8-Egfp/+; dsx-GAL4/+* |
| Supp. 2B | *dve-GAL4, nsyb-GAL80* | *w; UAS-cd8-Egfp/+; dve-GAL4, nsyb-GAL80/+* |
| Supp. 2C | *dve-GAL4, nsyb-GAL80* | *w, UAS-Dcr2; +/+; dve-GAL4, nsyb-GAL80/+* |
|  | *dve-GAL4, nsyb-GAL80, cornutus-IR, 1* | *w; P{GD15054}v29663/+; dve-GAL4, nsyb-GAL80/+* |
|  | *dve-GAL4, nsyb-GAL80, cornutus-IR, 2* | *w; P{KK109689}VIE-260B/+; dve-GAL4, nsyb-GAL80/+* |

